# Visual periodicity reveals distinct attentional signatures for face and non-face categories

**DOI:** 10.1101/2023.06.22.546207

**Authors:** Genevieve L. Quek, Adélaïde de Heering

**Affiliations:** The MARCS Institute for Brain, Behaviour and Development, Western Sydney University, Sydney, Australia; Unité de Recherche en Neurosciences Cognitives (UNESCOG), Center for Research in Cognition & Neurosciences (CRCN), Université libre de Bruxelles (ULB), Brussels, Belgium

**Keywords:** face perception, selective attention, EEG, frequency tagging, object recognition

## Abstract

Observers can selectively deploy attention to regions of space, moments in time, specific visual features, individual objects, and even specific high-level categories – for example, when keeping an eye out for dogs while jogging. Here we exploited visual periodicity to examine how category-based attention differentially modulates selective neural processing of face and non-face categories. We combined electroencephalography (EEG) with a novel frequency-tagging paradigm capable of capturing selective neural responses for multiple visual categories contained within the same rapid image stream (faces/birds in Exp 1; houses/birds in Exp 2). We found that the pattern of attentional enhancement and suppression for face-selective processing is unique compared to other object categories: Where attending to non-face objects strongly enhances their selective neural signals during a later stage of processing (300-500ms), attentional enhancement of face-selective processing is both earlier and comparatively more modest. Moreover, only the selective neural response for faces appears to be actively suppressed by attending towards an alternate visual category. These results underscore the special status that faces hold within the human visual system, and highlight the utility of visual periodicity as a powerful tool for indexing selective neural processing of multiple visual categories contained within the same image sequence.

## INTRODUCTION

With sensory input surging through the human visual system during our every waking moment, effective behaviour demands a mechanism for prioritizing relevant and important information within this immense data stream (Summerfield and Egner 2009). Selective attention is the process by which the brain filters incoming information on the basis of both bottom-up (i.e., visual salience) and top-down factors (i.e., relevance to current goals) (Johnston and Dark 1986; Corbetta and Shulman 2002), prioritizing the selected input for subsequent higher processing (Allport 1987). Where bottom-up attention is deployed rapidly and reflexively in response to salient visual events/features/stimuli (e.g., a peripheral flash) (Posner 1980; Müller and Rabbitt 1989), top-down attention is comparatively slower and requires more volitional effort on our part – hence the idiom “to pay attention” (Baluch and Itti 2011). At the same time, prioritising what is most relevant to our real-world goals is often controlled by a dynamic interplay between bottom-up and top-down factors, wherein feature-based attentional sets (Maunsell and Treue 2006) are subject to influence by expectations and knowledge about the world (Gayet and Peelen 2022; Yeh and Peelen 2022). Thus, if you lose your dog at the park, you might search for black, smallish objects, preferentially deploying this search template within the region of space where the target is most probable (e.g., on the grass, rather than in the lake).

In addition to individual visual features (Maunsell and Treue 2006) and their conjunctions (Egeth et al. 1984), visual selection can also be performed at the level of object categories (Nako et al. 2014; Stein and Peelen 2017; Battistoni et al. 2018; Quek, Nemrodov, et al. 2018; Störmer et al. 2019; Addleman et al. 2022). One category that holds unique importance for humans is faces, which convey a great deal of meaningful information that guides effective behaviour in social environments (e.g., age, sex, identity, emotion, and more). In particular, the human visual system appears remarkable adept at what has been referred to as *generic face recognition* (de Heering and Rossion 2015; Jacques et al. 2016; Quek, Liu-Shuang, et al. 2018; Quek et al. 2021), that is, categorising a face *as a face.* Subtended by occipitotemporal regions (Bentin et al. 1996; Kanwisher et al. 1997; Liu et al. 2002; Le Grand et al. 2003), this ability develops remarkably quickly during infancy (de Heering and Rossion 2015; Rekow et al. 2021) and arises based on very coarse visual input (Crouzet et al. 2010; Crouzet and Thorpe 2011; Quek, Liu-Shuang*, et al.* 2018; Quek *et al*. 2021). In fact, we are so good at detecting faces that we sometimes see a face when, in fact, there is no face present, as in the case of face pareidolia (Wardle et al. 2020; Rekow et al. 2022).

Given its status as a uniquely robust brain function, it is somewhat surprising to find that the interplay between face processing and selective attention has historically been examined under sparse and simplified conditions (e.g., binary classification of a spatiotemporally isolated stimulus, e.g., face or house). Moreover, attention to faces has largely been operationalised as a unitary construct, in which faces function as either the actively attended or explicitly ignored category (Wojciulik et al. 1998; Downing et al. 2001; Vuilleumier et al. 2001; Holmes et al. 2003; Engell and McCarthy 2010). Rapid visual presentation designs that exploit periodicity (Liu-Shuang et al. 2014; de Heering and Rossion 2015; Rossion et al. 2015) break with this standard, offering a way to quantify attentional effects for face-selective processing under high-competition conditions (i.e., distinguishing naturalistic images of faces amid a wide variety of other object categories). This approach has recently revealed the degree to which face-selective signals are respectively enhanced and suppressed (relative to a neutral baseline) by actively attending to faces as a category, or else towards another high-level category within the sequence (e.g., guitars, Quek, Nemrodov, et al. 2018). However, since Quek et al. (2018) only quantified the selective response to faces amid objects, it remains an open question as to how attentional selectivity at the level of object categories is achieved for non-face stimuli. To what extent are selective neural signals for an attended non-face category amplified relative to baseline? And equally, to what extent are task-irrelevant non-face categories are suppressed? Here we aimed to understand whether the relative contribution of enhancement and suppression to category-based attentional selectivity is similar across face and non-face categories. On one hand, perhaps when visual competition is high, task-relevance bestows a similar attentional benefit on selective visual processing regardless of which specific high-level category is the focus of current task demands. On the other hand, as a high-priority stimulus within the human visual system, faces could well be expected to exhibit a unique attentional profile compared with other visual categories (Lueschow et al. 2004; Reddy et al. 2004; Reddy et al. 2006; Finkbeiner and Palermo 2009).

To contrast how category-based attention modulates selective visual processing of face and non-face stimuli in the same group of observers, we adapted the fast periodic visual stimulation paradigm (de Heering and Rossion 2015; Rossion *et al*. 2015; Retter and Rossion 2016) to index selective neural signals associated with <u>multiple</u> high-level visual categories contained within the same image sequence. Observers saw a rapid sequence of various object images appearing at a strict 6 Hz presentation rate, with two critical categories of interest interlaced at distinct periodicities within the image stream (e.g., 1 in 5 images, 1 in 4 images). Coupled with a neurophysiological measure such as electroencephalography (EEG), this manipulation effectively tags the selective neural response to each critical category with its own pre-specified stimulation frequency (i.e., 1.2 Hz & 1.5 Hz). This ‘interlaced periodicities’ approach is inspired by, but distinct from, the so-called ‘steady-state’ or ‘temporal-tagging’ method wherein two concurrently presented stimuli (or even two parts of a single stimulus, e.g., Boremanse et al. 2013) flicker at distinct frequencies, which has been used effectively to study the influence of attention on visual processing in prior studies (Müller et al. 2006; Appelbaum and Norcia 2009; Jaeger et al. 2018; Brummerloh et al. 2019).

In Experiment 1, we used this approach to derive separable neural indices of face-selective visual processing and bird-selective visual processing. Manipulating the focus of category-based attention across blocks, we quantified both category-selective signals in the context of an *attend-Faces* task and an *attend-Birds* task, and contrast these against the same signals measured during a neutral attentional baseline condition in which observers monitored the fixation cross for colour changes. Note that birds were not intended as a specific control category for faces, but rather as another animate category for which attention could be expected to modulate the visual response. Moreover, birds have been used effectively in other periodicity-based studies (Hagen and Tanaka 2019). Experiment 2 was near identical, save that houses replaced faces as the second critical category. To anticipate our results, we observed distinct profiles of attentional enhancement and suppression for selective visual processing associated with individual high-level categories (i.e., faces/birds/houses). In Exp 1, attending to faces enhanced their category-selective neural response to a much lesser extent than was evident for birds. Conversely, attentional suppression induced by attending to the alternate critical category was pronounced for face-selective processing, but non-existent for bird-selective processing. In Exp 2, we confirmed that this pattern across face and non-face categories did not arise as an artefact of the specific stimulation frequencies by substituting houses for faces. Temporal analyses revealed distinct time courses of attentional effects across the different categories; where enhancement of face-selective processing was most evident before 200 ms, the bird- and house-selective waveforms were not facilitated by attention until much later (i.e., 300-500ms).

## MATERIALS & METHODS (Experiments 1 & 2)

### Participants

The study was carried out in accordance with the guidelines and regulations of the Research Ethics Board of the department of Psychology of the Université libre de Bruxelles (Belgium) (Reference - B406201734083). We tested two different groups of healthy adult volunteers who reported no psychiatric or neurological disorders. Twenty-one participants took part in Exp 1 in exchange for financial compensation, and 23 participants took part in Exp 2 in exchange for course credit. All were right-handed, reported having normal or corrected-to-normal vision, and gave written informed consent prior to the start of the experiment. Several participants’ data were discarded due to excessive noise (1 from Exp 1, 3 from Exp 2), with the final sample in each experiment consisting of 20 individuals (Exp 1 = 3 males; mean age 20±1.8 (SD) years; Exp 2 = 5 males; mean age 24±4 (SD) years).

### Stimuli

Stimuli were a previously described set of 200 x 200-pixel colour images (de Heering and Rossion 2015; Rossion *et al*. 2015; Jacques *et al*. 2016) that included 48 faces, 24 birds, 48 houses, and a further 176 images drawn from various categories (e.g., chairs, guitars, flowers, animals…). Image subjects were left embedded in their original backgrounds, yielding an image set that varied widely in colour composition, viewpoint, lighting condition, etc. (see Figure 1A). Each image subtended 6.86 x 6.86 degrees of visual angle at a viewing distance of 50 cm.

**Figure 1.**
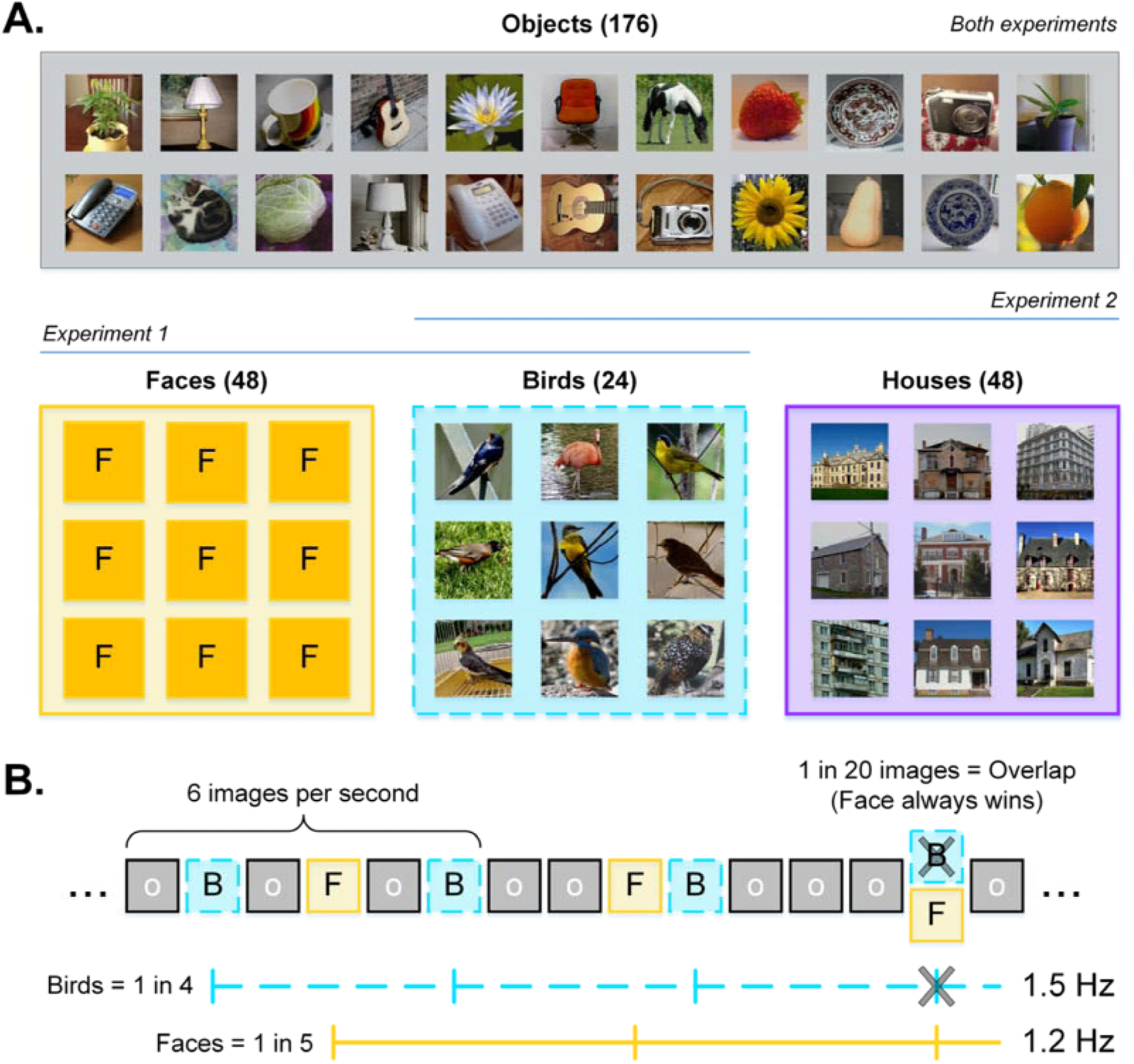
**A)** Examples of category exemplars used in Exp 1 (objects, faces, birds) and Exp 2 (objects, houses, birds). Instances of guitars (additional targets in Exp 2) are given in the Object panel. **B)** Schematic of the ‘interlaced’ sequence in Exp 1. Objects appeared at a rate of 6 Hz, with bird and face exemplars appearing as every 4^th^ and 5^th^ image, respectively. Where these 1.5 and 1.2 Hz frequencies overlapped every 20 images, a face image always took precedence (i.e., effectively skipping a bird presentation), allowing for exactly 48 bird and 48 face instances in each 40 second sequence. Sequence structure in Exp 2 was identical, save that house images replaced the face images.

### Apparatus

The experiments took place in a darkened, sound-attenuated room, where participants sat before a Lenovo monitor (1920×1080; 60Hz refresh rate) that displayed a uniform grey background throughout the recording session. Image sequence presentation was programmed in Psychtoolbox (MATLAB, The MathWorks). We recorded scalp electrical activity using a 64-channel BioSemi ActiveTwo system (Amsterdam, Netherlands) with a sample rate of 1024Hz. During recording setup, we held individual electrode offsets below ±30 μV by softly abrading the participant’s scalp with a blunt plastic needle and insulating the electrode tip with gel. We monitored eye movements in Exp 1 via 4 external electrodes (one placed at the outer canthi of each eye, and one placed immediately above and below the right orbit). In Exp 2, we achieved the same purpose by visualising eye movement behaviour via an external camera.

### Design (Experiments 1 & 2)

#### Sequence Composition

Each sequence began with a “ready?” message. Once the experimenter confirmed that the EEG trace was artefact-free, the participant pressed the spacebar to launch the sequence. A black fixation cross appeared in the centre of the screen for a random interval between 1 to 5 seconds, after which they saw a rapid stream of various objects (e.g., vehicles, animals, trees, structures, etc.) appearing at a strict periodic rate of 6 images per second. To achieve this 6 Hz stimulation, we sinusoidally modulated the contrast of each image from 0-100-0% over 167ms (i.e., 10 screen refreshes). Within the image stream, two critical categories (birds/faces for Exp 1; birds/houses for Exp 2) were interlaced at distinct embedded periodicities: A bird appeared as every 4^th^ image, and a face (Exp 1) or house (Exp 2) as every 5^th^ image (Figure 1B). We marked the onsets for object, bird, and face/house stimuli in the EEG trace using distinct numeric triggers. As detailed elsewhere (de Heering and Rossion 2015; Rossion *et al*. 2015; Jacques *et al*. 2016; Quek and Rossion 2017), this visual stimulation reliably elicits neural responses at the exact frequencies of stimulation, with a general visual response arising at the image presentation rate (i.e., 6 Hz), and category-selective responses at the embedded periodicities. Importantly for our purposes, since the critical categories here appear at distinct periodic intervals in the sequence (i.e., 1/4 & 1/5 image respectively), their differential neural signatures are readily dissociable at 1.2 Hz for faces/houses and 1.5 Hz for birds. Where these frequencies overlapped (every 20 stimuli, see Figure 1B), the face (Exp 1) or house (Exp 2) was always presented, effectively skipping a bird instance. This served to equalize the total number of faces/houses and birds that appeared in each sequence: 48 occurrences of each critical stimulus type within each 40 second sequence. To encourage fixation and attention to the images, the fixation cross remained superimposed on the images throughout the entire sequence. To minimize blinks and artefacts elicited by the sudden appearance/disappearance of flickering stimuli, each 40-second sequence was preceded and followed by a transition period during which the maximum contrast of each image progressively ramped up or down (total sequence time = 43.33 seconds). These fade-in / fade-out periods were not included in analysis.

#### Attentional Manipulation

Participants in Exp 1 saw three blocks of 15 sequences that were strictly identical in visual content, varying only in terms of the task (order counterbalanced across participants). In the *attend-Cross* block, participants had to press the spacebar whenever the overlying fixation cross changed colour from black to blue (200ms change duration, 8 random occurrences per sequence). This orthogonal change detection task is the paradigm standard (Liu-Shuang *et al*. 2014; Rossion *et al*. 2015; Jacques *et al*. 2016; Quek and Rossion 2017; Quek, Nemrodov*, et al.* 2018) and was used as a baseline condition that encouraged a constant level of attention to the images themselves (which were irrelevant to the colour-change task). In the *attend-Birds* block, participants responded whenever a bird appeared in the image stream (i.e., 1 in 4 images), and in the *attend-Faces* block, whenever a face appeared (i.e., 1 in 5 images). Supplemental Figure S5 presents the response time (RT) data associated with this task, with the caveat that the periodic nature of targets might somewhat compromise its utility as a meaningful behavioural index of face/bird detection. To take account of the possibility that motor responses on the behavioural task could contribute to the neural response we index at the category-specific frequencies, we also inspected Lateralised Readiness Potentials (LRPs) – these results are presented in Supplemental Figure S2.

The structure of Exp 2 was near-identical, save that the *attend-Faces* block became an *attend-Houses* block, with images of houses replacing the faces to become the second critical category. In Exp 2, we included a fourth block of sequences in which the task was to attend to a target category that appeared non-periodically throughout the sequence (guitars). The *attend-Guitars* block was always presented last, with the first three task-blocks counterbalanced as in Exp 1. The total number of trials in Exp 1 and Exp 2 was thus 45 and 60 respectively. The 15 sequences per block lasted ∼10 minutes. Rest breaks were self-paced.

### EEG Analyses

#### EEG Pre-processing

We analysed EEG data in Letswave6 (https://www.letswave.org/) running in MATLAB (MathWorks). After importing each participant’s raw continuous recording, we applied a band-pass filter (0.1-100Hz) before downsampling the data to 250Hz for faster handling and storage. We segmented 48-second epochs corresponding to each sequence, starting from 2 seconds before the onset of the sequence and lasting until 46 seconds. Noisy electrodes (identified by eye) were linearly interpolated using two immediately adjacent clean channels (no more than 4 channels interpolated per participant). We re-referenced all scalp channels to the average of all 64 channels before cropping each epoch to exactly 40 seconds, corresponding to the sequence proper (i.e., excluding the fade in/fade out period). This produced 15 epochs per task condition, per participant.

#### Frequency Domain Analyses

### Response Significance & Harmonic Selection

To determine the harmonic range over which to quantify *i)* the common visual response and *ii)* the category-selective responses in Exp 1, we obtained the grand mean amplitude spectrum by averaging epochs across all participants, conditions, and scalp channels and subjecting the resulting waveform to Fast Fourier Transformation (FFT). We computed a *z*-score for each frequency bin (z = (x - µ)/a), where x is the amplitude at a frequency bin of interest, µ is the mean amplitude of the 14 bins surrounding x (i.e., 7 bins either side, excluding the immediately adjacent bin), and a is the standard deviation across that same 14 bin range. With our frequency resolution of 0.025 (i.e., 1/40 seconds), this 14-bin range ensured that noise estimates were based on a similar frequency range as in Quek et al. (2018) (i.e., 0.35 Hz *vs.* 0.36 Hz). We inspected *z* scores on the specific harmonic frequencies of the image presentation rate (6 Hz), bird presentation rate (1.2 Hz), and face presentation rate (1.5 Hz). Continuous harmonics at which *z* > 3.1 (i.e., *p* < .001, one-tailed, signal>noise) were included for response quantification; a conservative criterion we have imposed in several previous periodicity-based studies (Quek and Rossion 2017; Quek, Nemrodov*, et al.* 2018). For Exp 1, this identified the first 6 harmonics of the image presentation rate (i.e., Common Visual response = 6, 12, 18, 24, 30, & 36 Hz), the first 11 selective harmonics of the 1.5 Hz bird rate (i.e., bird-selective response = 1.5, 3, 4.5, 7.5, 9, 10.5, 13.5, 15, 16.5, 19.5, 21 Hz) and the first 16 harmonics of the 1.2 Hz face rate (i.e., face-selective response = 1.2, 2.4, 3.6, 4.8, 7.2, 8.4, 9.6, 10.8, 13.2, 14.4, 15.6, 16.8, 19.2, 20.4, 21.6, & 22.8 Hz). Note that harmonics of 6Hz were never included in quantification ranges for the category-selective responses. Identified harmonic ranges in Exp 2 were near-identical: Common Visual Response = 6-36Hz, Bird-selective response = 1.5-16.5 Hz (i.e., two fewer harmonics than in Exp 1), House-selective response = 1.2-21.6 Hz (one fewer harmonic than in Exp 1). Since amplitudes at these upper harmonics are very small (see Figure 3A/5A), their inclusion in the quantification stage does not influence the overall pattern of results. Thus, for consistency’s sake, we took the harmonic ranges identified in Exp 1 as the standard for both experiments.

### Response Quantification

To quantify the common visual response, we averaged the 15 epochs per condition for each participant and subjected these conditional means to FFT. At each frequency bin, we applied a baseline-correction by subtracting the mean amplitude of the local noise range (i.e., 14 bins as described above for Z scores). We then summed baseline-corrected amplitudes across the identified 6-36Hz harmonic range to produce a final value that was subjected to statistical analysis. We quantified the bird- and face/house-selective signals for each individual participant in the same way, save that we performed an additional cropping step prior to FFT: First, we cropped the conditional means to 39.34 seconds (9834 bins), reflecting an integer number of cycles of the bird presentation frequency at 1.4999 Hz. Separately, we cropped the conditional means to 39.17 seconds (9792 bins), reflecting an integer number of cycles of the face presentation frequency at 1.1999 Hz. These different crop lengths ensured that the frequency resolutions of the subsequent FFT amplitude spectra were appropriate for the relevant signal of interest (i.e., with a bin centred exactly on the fundamental frequency under inspection). We then performed a baseline-correction using the local noise range (see above), before computing category-selective summary scores by summing across the relevant harmonic ranges (birds: 1.5-21 Hz, faces = 1.2-22.8 Hz, excluding harmonics of 6 Hz). Conditional data for each signal type was then relabelled to denote the relationship between the focus of the participant’s task in that block and the signal type. For the face-selective signal, the *attend-Faces* block was relabelled **Attend Towards**, and the *attend-Birds* block relabelled **Attend Away**. The reverse was true for the bird-selective signal: Here, *attend-Faces* became **Attend Away** and *attend-Birds* became **Attend Towards**. In both cases, the *attend-Cross* block was relabelled as **Baseline**. Individual participant summed amplitude values in these relabelled conditions were subjected to statistical analysis. As a final step, we followed Quek et al. (2018) by computing indices of attentional enhancement and attentional suppression for each category selective signal (Enhancement = Attend Towards minus Baseline; Suppression = Attend Away minus Baseline). In Exp 2, we calculated the indices twice: once using *attend-Cross* as the baseline and once using *attend-Guitar*.

### Regions of Interest

We defined two occipitotemporal (OT) regions-of-interest or ROIs (*Left OT*: PO3, P7, PO7, P9, O1; *Right OT*: PO4, P8, PO8, P10, O2) that have been shown to capture neural responses associated with high-level visual discrimination (Jacques *et al*. 2016; Quek, Liu-Shuang*, et al.* 2018; Quek, Nemrodov*, et al.* 2018; Hagen and Tanaka 2019; Quek *et al*. 2021). Category-selective responses were examined in these bilateral OT ROIs. We also defined an occipital ROI (O1, O2, Oz) where the 6 Hz response was expected to be maximal; we contrasted the common visual response across all three ROIs.

### Statistical Analyses

Signals were compared across conditions using i) repeated measures ANOVAs, applying a Greenhouse-Geisser correction wherever the assumption of sphericity was violated, ii) paired *t*-tests with Bonferroni corrections, and iii) Bayesian paired *t*-tests. All attentional indices were evaluated against zero using one-sample *t*-tests with a Bonferroni correction.

#### Time Domain Analyses

To inspect each participant’s category-selective responses in the time-domain, we subjected the preprocessed epochs to a 4^th^ order Butterworth low-pass filter with a cutoff of 30Hz. We then removed the common visual response at 6 Hz and harmonics using an FFT multi-notch filter (width = 3 bins) centred on each of the first 6 harmonics of the image presentation frequency (slope cutoff width = 0.0750Hz). We segmented an 833.33 ms window around each Face/House or Bird occurrence starting 167 ms before stimulus onset, discarding the final epoch of each stimulus type in each sequence since it overlapped the fade-out period (totalling 705 bird epochs and 525 face/house epochs). We then averaged sequences within each condition for each participant and performed a baseline correction by subtracting the mean amplitude in the -167ms to 0ms time-window from the entire epoch. Finally, we averaged the baseline-corrected conditional means across participants to produce group means for visualization. Condition relabelling was applied as in the frequency-domain analysis (see above). We assessed the presence of enhancement and suppression at each timepoint via paired Bayesian *t*-tests (two tailed) for the relevant condition contrasts (i.e., Baseline *vs.* Attend Towards, and Baseline *vs.* Attend Away). Responses were inspected in the same left and right OT ROIs as described above for the frequency-domain analyses.

## RESULTS

### EXPERIMENT 1 RESULTS

#### Frequency Domain

Figure 2A presents the grand averaged amplitude spectrum for Exp 1. As is standard in SS-EP designs (Liu-Shuang *et al*. 2014; Rossion 2014; Jacques *et al*. 2016; Quek and Rossion 2017; Quek, Nemrodov*, et al.* 2018), we observed a strong response at the image presentation frequency (i.e., 6Hz and harmonics), which is understood to capture aspects of visual processing shared by all image categories contained within the sequence. Moreover, we found clear evidence for selective visual processing associated with both birds and faces, with significant signal at their respective presentation frequencies and harmonics (shaded bars in Figure 2A).

**Figure 2.**
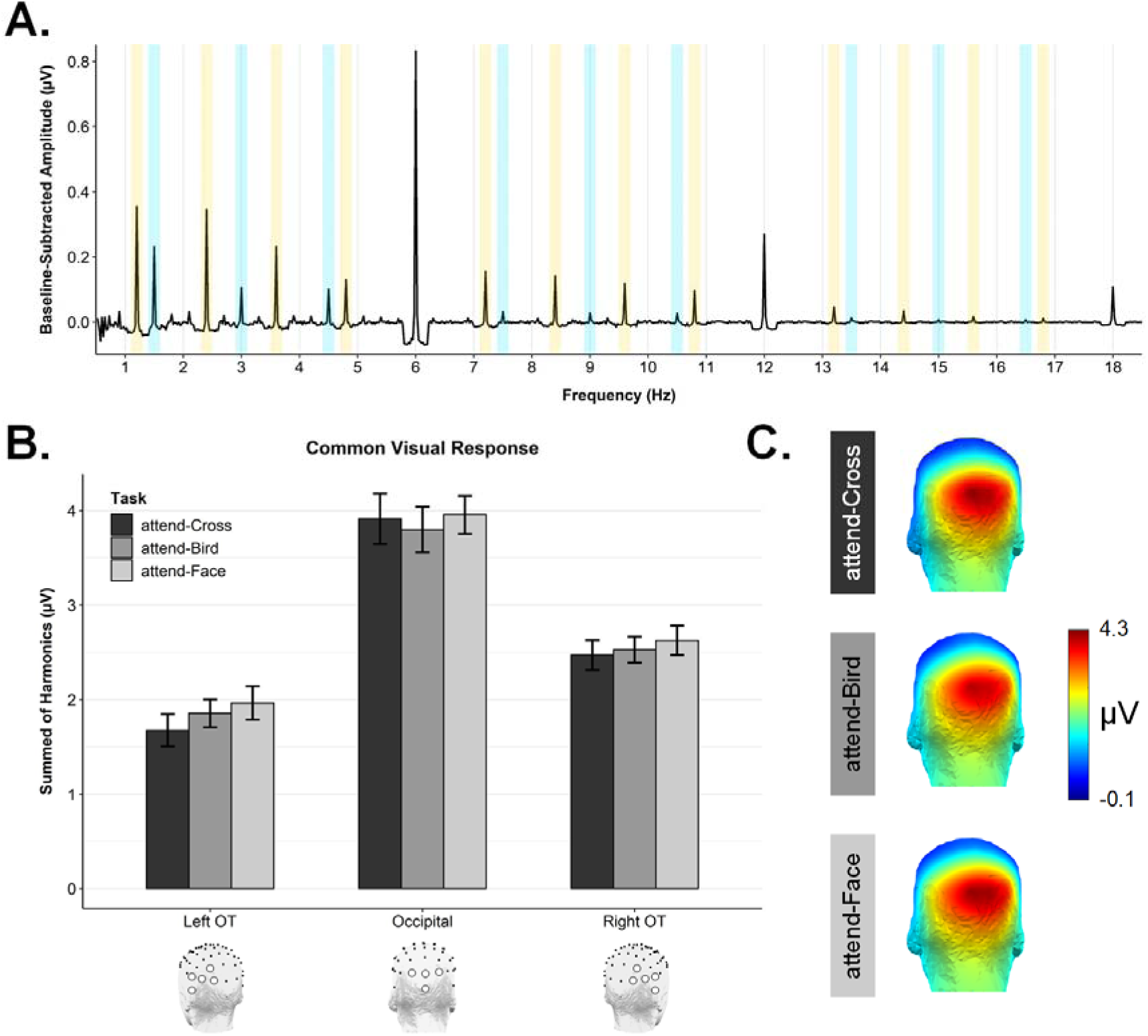
**A)** The grand averaged amplitude spectrum for Exp 1, shown up to 18 Hz for visualisation purposes. Strong and significant responses were evident at the image presentation frequency (i.e., 6 Hz & harmonics), face-selective frequency (1.2 Hz & harmonics, yellow shaded regions), and bird-selective frequency (1.5 Hz & harmonics, blue shaded regions). **B)** The Common Visual Response for Exp 1 (i.e., sum of 6, 12, 18, 24, 30, & 36 Hz), shown as a function of Task and ROI. **C)** Scalp topographies for the Exp 1 common visual response in each Task condition.

**Figure 3.**
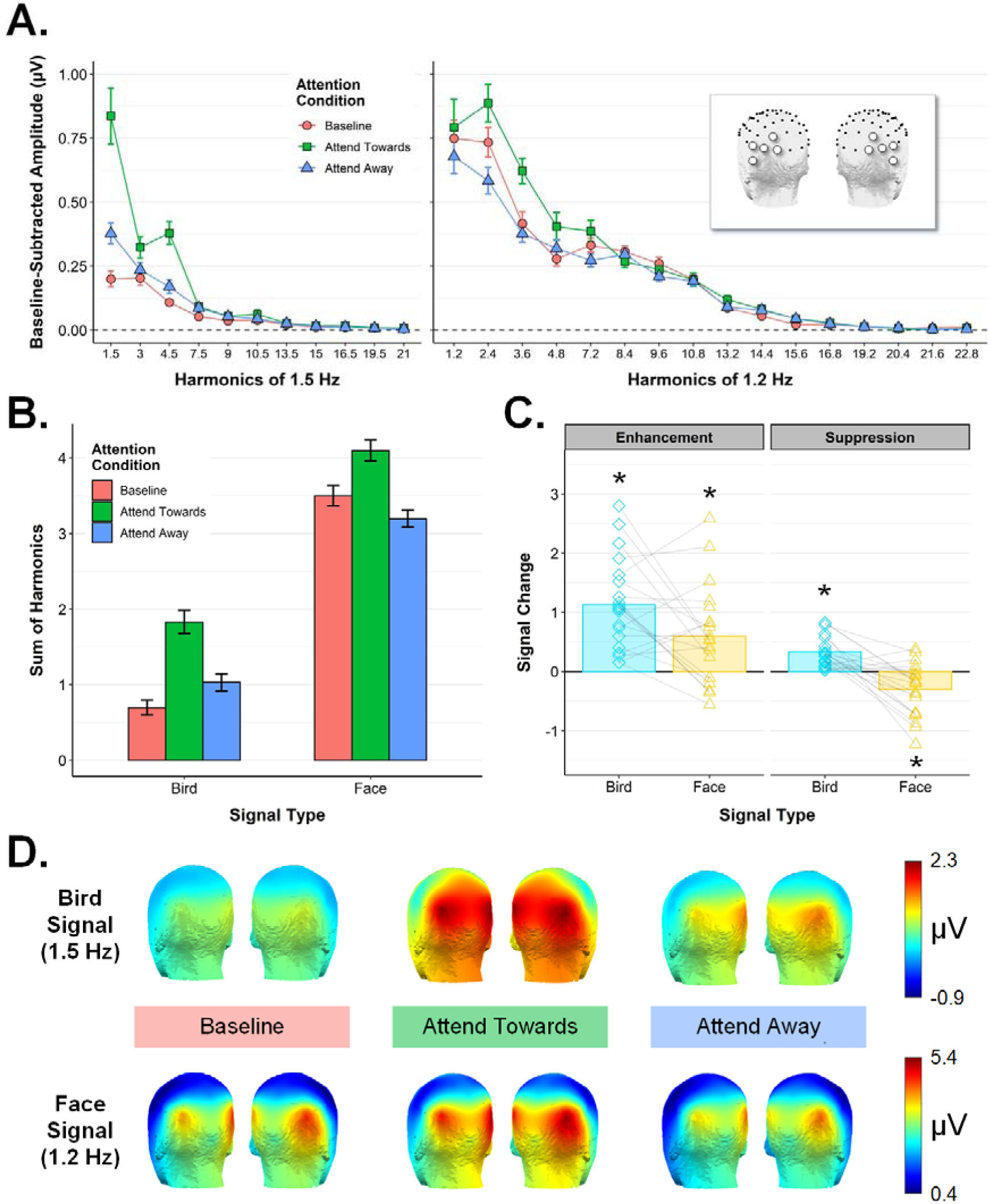
Exp 1 frequency domain results. **A)** Conditional mean amplitudes at each harmonic frequency included in the quantification of the bird-selective response (left panel) and face-selective response (right panel), averaged across the left/right ROIs (see inset). All error bars are within-subjects standard error. **B)** The quantified bird- and face-selective responses (i.e., amplitudes summed across the harmonic ranges indicated in A), averaged across the left/right ROIs. **C)** Indices of Enhancement (Attend Towards – Baseline) and Suppression (Attend Away – Baseline) for the bird- and face-selective responses. Overlaid points are individual participants. \**p* < .05, Bonferroni-corrected, one sample *t*-test against zero). **D)** Conditional mean scalp topographies for the bird- (top row) and face-selective responses (bottom row) in Exp 1. Amplitude ranges are fixed across Attention Condition for each signal type. Individual participant topographies for each condition are given in Supplemental Figure S1A.

We subjected the common visual response values to a 3×3 repeated measures ANOVA with the factors *ROI* (left OT, right OT, occipital), and *Task* (attend-Cross, attend-Birds, attend-Faces). There was a significant main effect of *ROI*, *F*(1.47,28.02) = 23.92, *p <* .00001, ^2^ = .311, with notably larger amplitudes, as expected, in the occipital ROI (*M* = 3.89, *SD* = 1.83) than in the left OT (*M* = 1.83, *SD* = 0.68) or right OT (*M* = 2.54, *SD* = 0.98) ROIs, see Figure 2B. Figure 2C shows the quantified common visual response in each condition is centred over the occipital pole. In contrast, there was no significant influence of *Task* on the common visual response, *F*(2,38) = 1.61, *p =* .213, □^2^ = .003, nor was there a significant interaction between *ROI* and *Task*, *F*(2.32,44.05) = 1.36, *p =* .269, □^2^ = .002. The consistent common visual response across Task conditions suggests that participants’ overall level of arousal did not differ as a function of the specific attentional task.

Next, we examined the summarised face-selective and bird-selective responses in each attention condition (see scalp topographies in Figure 3D) via a 3-way repeated measures ANOVA with the factors *Signal Type* (face, bird), *Attention Condition* (Baseline, Attend Towards, Attend Away), and *ROI* (left OT, right OT). We found a main effect of *Signal Type*, *F*(1,19) = 169.39, *p* < .00001, □^2^ = .613, reflecting, as expected, larger amplitudes for the face-selective response (*M* = 3.6, *SD* = 0.97) than the bird-selective response (*M* = 1.18, *SD* = 0.48) (see Figure 3B). There was a main effect of ROI, *F*(1,19) = 6.58, *p* < .05, □^2^ = .042, with larger amplitudes in the right OT ROI (*M* = 2.59, *SD* = 0.93) than in the left OT ROI (*M* = 2.19, *SD* = 0.47). There was also a main effect of *Attention Condition, F*(1.23,23.32) = 35.56, *p* < .00001, □^2^ = .151, as well as a significant *Signal Type* x *Attention Condition* interaction*, F*(1.51,28.64) = 7.02, *p* = .006, □^2^ = .021. *ROI* did not interact significantly with either *Signal Type*, *F*(1,19) = 3.43, *p* = .079, □^2^ = .012, or *Attention Condition*, *F*(1.33,25.32) = 0.22, *p* = .713, □^2^= .000. The 3-way interaction did not reach significance either, *F*(2,38) = 0.94, *p* = .398, □^2^ = .001.

Figure 3C shows indices of attentional enhancement (i.e., Attend Towards – Baseline) and suppression (i.e., Attend Away – Baseline) for each signal type. As difference scores calculated within signal type, these indices are not affected by the higher overall amplitude of the face-selective response compared to the bird-selective response. After confirming that each index departed significantly from zero (all one-sample *t*-test *p* values < .05, Bonferroni-corrected), we used a one-way ANOVA to ask whether each metric differed as a function of *Signal Type*. As Figure 3C shows, enhancement was significantly stronger for the bird signal (*M* = 1.13, *SD* = 0.77) compared to the face signal (*M* = 0.60, *SD* = 0.82), *F*(1,19) = 5.99, *p* = .024, □^2^ = .106. In contrast, directing attention to the alternate visual category served to suppress the face-selective signal (*M* = -0.30, *SD* = 0.44) to a greater extent than the bird-selective signal (*M* = 0.34, *SD* = 0.26), *F*(1,19) = 26.82, *p* < .0001, □^2^= .446. In fact, suppression scores for the bird signal were positive on average, suggesting that attending to faces within the image stream did not suppress the evoked response to birds at all, but in fact slightly facilitated this category-selective response as compared to the *attend-Cross* baseline.

#### Time Domain

To understand how category-based attention modulated the bird- and face-selective signals over time, we inspected stimulus-locked responses to birds and faces in each attention condition after notch-filtering the response at 6 Hz and harmonics. The selective waveform for faces (Figure 4A) was highly consistent with what has been reported previously using periodicity-based paradigms (Rossion *et al*. 2015; Jacques *et al*. 2016; Retter and Rossion 2016; Quek and Rossion 2017), evolving over some 600ms following stimulus onset^1^. The selective response elicited by birds presented amid various other natural object categories (Figure 4B) is a novel report that has not previously been characterized. Bayesian paired *t*-tests contrasting amplitudes for the Attend Towards and Baseline conditions at each timepoint revealed markedly different patterns of attentional enhancement for face- and bird-selective processing: Although both signals showed a facilitation effect during the early stages of visual processing (with slightly later facilitation evident for birds as compared to faces), the bird-selective response was most strongly enhanced during a later time window (∼300-500ms). Notably, this late facilitation was entirely absent for the face-selective response. Since we wondered how this timecourse might overlap with action preparation signals, we also inspected lateralised readiness potentials (LRPs) in each condition by subtracting electrode C4 amplitudes from C3 amplitudes, separately for the face and bird locked waveforms. We were specifically interested in whether LRPs would be strongly evident in the Attend Towards condition (which requires a behavioural response each time that attended category appears), however this was not the case (see Figure S2 in Supplemental Materials).

**Figure 4.**
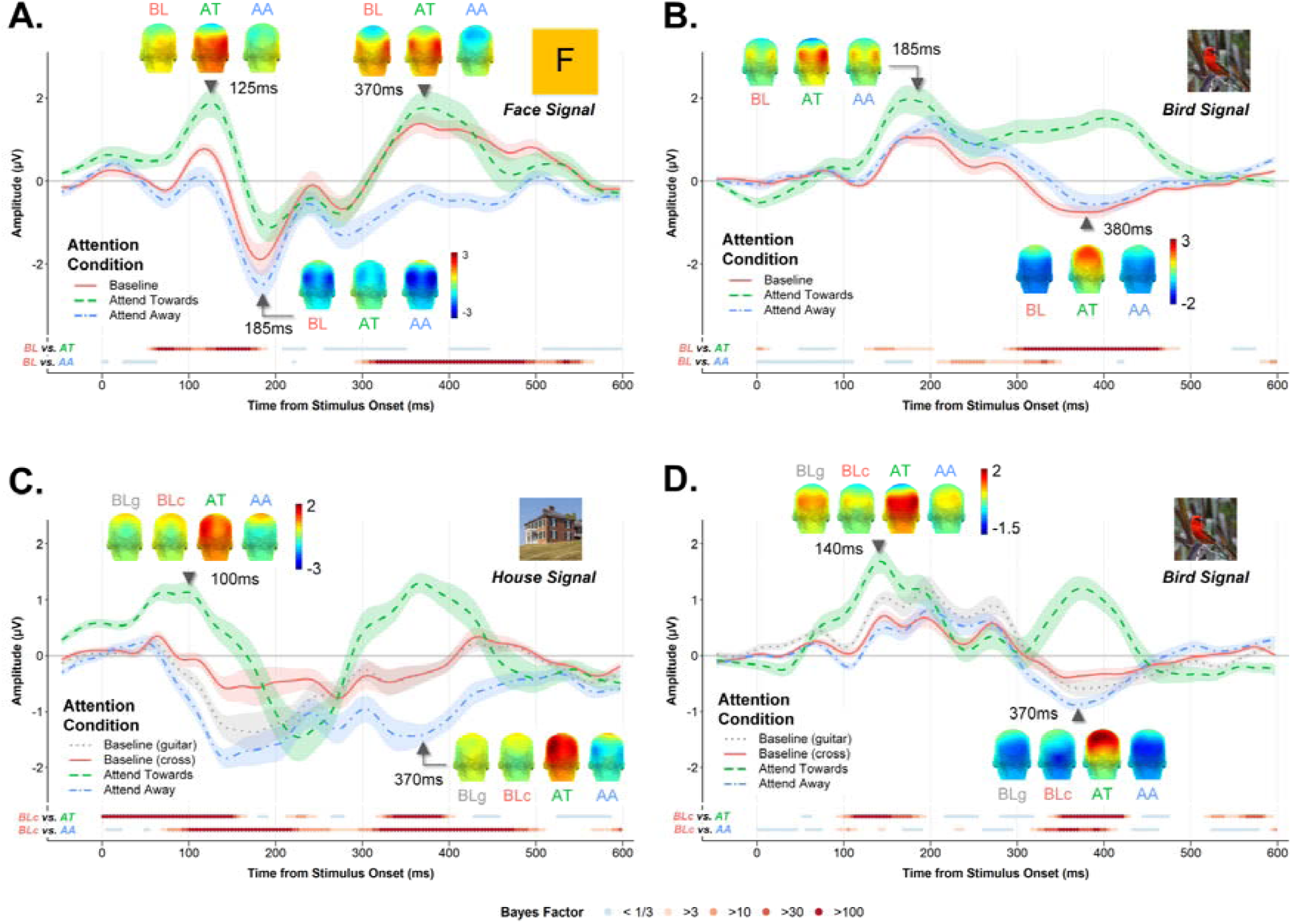
Category-selective responses in Exps 1 (A & B) and 2 (C & D), shown as a function of time from stimulus onset for each attention condition. **A)** The face-selective and **B)** bird-selective responses in Exp 1, averaged across the left and right OT ROIs. Shaded regions are within-subjects standard error. Coloured points below each plot reflect Bayesian evidence at each timepoint for a difference between the Baseline (BL) & Attend Towards (AT) conditions and the Baseline (BL) & Attend Away (AA) conditions. Cool and warm colours denote evidence for the null and H_1_, respectively. Group-averaged headplots are given for select timepoints; amplitude ranges are fixed within signal type (i.e., same colourbar for all headplots for each signal). **C & D)** As above, but for the House-selective and Bird-selective signals in Exp 2. Category-selective waveforms separated by left and right OT ROIs are given in supplemental Figures S3 & S4.

Conversely, contrasting the Attend Away and Baseline conditions revealed that attending to Birds significantly dampened the amplitude of the face-selective response from ∼300ms until nearly 600ms post stimulus onset. This window of attentional suppression corresponds well with what Quek et al. (2018) observed for face-selective responses, although in our case the suppression effect endured for longer. Notably, the reverse effect was not evident: that is, attending to faces did not have a dampening effect on the bird-selective response, but in fact resulted in a response amplitude increase between 200-350ms.

### EXPERIMENT 2 RESULTS

#### Frequency Domain

Figure 5E shows the Exp 2 **common visual response** as a function of Task. A 3×4 repeated measures ANOVA with the factors *ROI* (left OT, right OT, occipital) and *Task* (attend-Cross, attend-Guitars, attend-Houses, attend-Birds) revealed significant main effects of *ROI*, *F*(2,38) = 12.78, *p* < .0001, □^2^ = .156, and *Task*, *F*(2.05,38.92) = 3.56, *p* = .037, □^2^ = .007. Since there was a significant interaction between these factors, *F*(3.01,57.14) = 3.29, *p* = .027, □^2^ = .003, we separated the occipital ROI from the OT ROIs for follow up. Importantly, we found that *Task* did not modulate common visual response magnitudes in the occipital ROI, *F*(3,57) = 2.50, *p* = .069, □^2^ = .006, where this general measure of arousal is strongest (Figure 5F). The 2×4 repeated measures ANOVA for the OT ROIs revealed a main effect of *ROI, F*(1,19) = 10.99, *p* = .004, □^2^ = .122 (stronger response in the right OT ROI, *M* = 2.8, *SD* = 1.52, compared to the left OT, *M* = 1.9, *SD* = 0.75). There was also a main effect of *Task, F*(1.99,37.89) = 4.95, *p* = .012, □^2^ = .012. As can be seen in Figure 5E, the attend-Cross task elicited the smallest common visual response in the OT ROIs, although all post-hoc contrasts between individual Task conditions were non-significant (pairwise t-test *p*-values all > .05). That the Common Visual Response did not vary greatly as a function of *Task* suggests that participants’ overall level of arousal did not vary greatly across the different attentional conditions.

**Figure 5.**
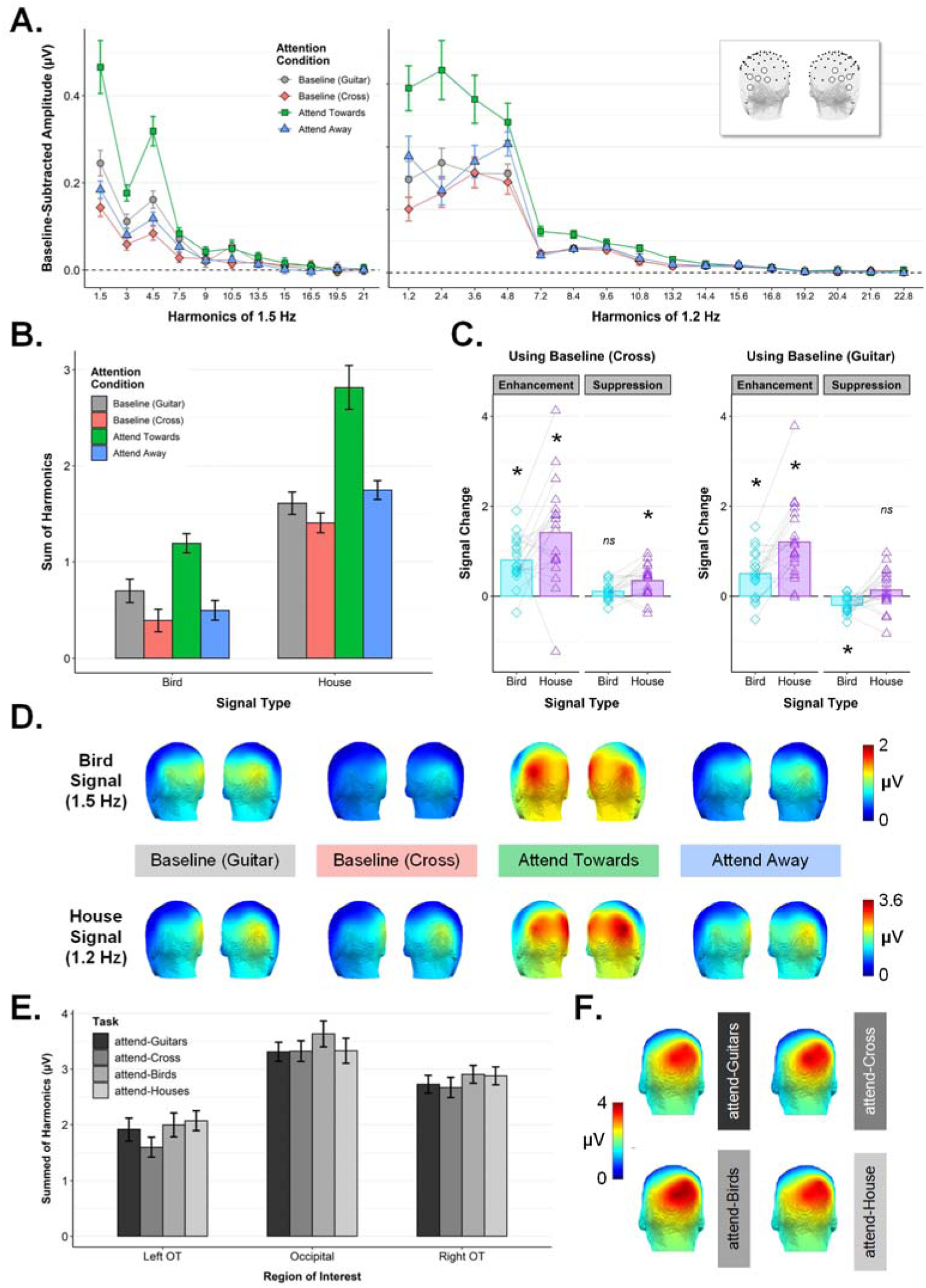
Exp 2 frequency domain results. **A)** Conditional mean amplitudes at each harmonic frequency of the bird-selective response (left panel) and house-selective response (right panel), averaged across the left/right ROIs (see inset). All error bars are within-subjects standard error. **B)** The quantified bird- and house-selective responses (i.e., amplitudes summed across the harmonic ranges indicated in A), averaged across the left/right ROIs. **C)** Indices of enhancement and suppression for the bird-selective (blue) and house-selective (purple) signals, calculated separately using *attend-Cross* as Baseline (left panel) and *attend-Guitar* as Baseline (right panel). Overlaid points are individual participants. (\**p* < .01, Bonferroni-corrected, one sample *t*-test against zero). **D)** Scalp topographies for the quantified bird-selective (top row) and house-selective response (bottom row) in each condition of Exp 2. Individual participant topographies are given in supplemental Figure S1B. **E.** The Common Visual Response shown as a function of Task and ROI. **F.** Scalp topographies for the common visual response in each Task condition.

Next, we inspected the category-selective responses across conditions (see scalp topographies in Figure 5D). We mirrored the structure of Exp 1 analyses by treating the *attend-Cross* condition as the relevant baseline. A 3-way repeated measures ANOVA with the factors *Signal Type* (house, bird), *Attention Condition* (Baseline, Attend Towards, Attend Away), and *ROI* (left OT, right OT) revealed a main effect of *Signal Type*, *F*(1,19) = 46.87, *p* < .00001, □^2^ = .318, reflecting larger amplitudes for the house-selective response (*M* = 1.99, *SD* = 1.01) compared to the bird-selective response (*M* = 0.69, *SD* = 0.25) (see Figure 5B). There was also a main effect of *Attention Condition*, *F*(1.19,22.59) = 44.25, *p* < .000001, ^2^ = .201, and a significant interaction between *Attention Condition* and *Signal Type, F*(1.12,21.19) = 4.61, *p* = .04, □^2^ = .017. The main effect of *ROI*, and all its interaction terms, did not reach significance (all *p-*values > .05). For indices of attentional enhancement and suppression calculated using the *attend-Cross* condition as the baseline (see Figure 5C, left panel), all indices departed significantly from zero (all one-sample *t*-test *p* values < .05, Bonferroni-corrected) save for the suppression score for the bird signal. We found that enhancement differed significantly depending on *Signal Type*, *F*(1,19) = 5.94, *p* = .025, □^2^ = .115, with stronger enhancement for the house-selective 1.2Hz signal (*M*_chang*e*_ = 1.41, *SD* = 1.13) compared to the bird-selective 1.5Hz signal (*M*_chang*e*_ = 0.8, *SD* = 0.5). Notably, this pattern across the two selective frequencies was the reverse of what we observed in Exp 1, *F*(1,38) = 11.55, *p* = .002, □^2^= .109, where we found stronger enhancement at 1.5 Hz compared to 1.2Hz. This is encouraging, since it suggests that the attentional differences observed for Birds and Faces in Exp 1 cannot be explained by the specific frequency of presentation assigned to each category. Suppression scores for both signal types were slightly positive, suggesting that neither category-selective signal was actively attenuated by attending to the other critical category, but instead slightly facilitated relative to the attend-Cross baseline (see Figure 5C, left panel). This somewhat paradoxical facilitation effect was greater for the house signal (*M*_chang*e*_ = 0.35, *SD* = 0.35) than for the bird signal (*M*_chang*e*_ = 0.1, *SD* = 0.18), *F*(1,19) = 13.05, *p* = .002, □^2^= .174, though only the former departed significantly from zero, *t*(19) = 4.43, *p*_bonf_ < .01. Notably, this pattern of numerically positive suppression for both signal types differed from what we observed in Exp 1, *F*(1,38) = 38.5, *p* < .000001, □^2^ = .322, where we found that attending to birds served to actively suppress face-selective processing, but attending to faces actually served to facilitate the bird-selective signal (see Figure 3C).

Next, we repeated the category-selective analysis using the *attend-Guitars* block as the neutral attentional baseline. We included this condition in Exp 2 to examine category-selective responses when attention was directed away from the critical categories, but still focused on the images themselves (by attending to the nonperodic instances of guitars in the sequence). We predicted this instruction would boost the bird- and house-selective signals relative to when attention was focused on the central fixation cross (i.e., grey *vs.* red bars in Figure 5B). A paired t-test confirmed this was the case, *t*(19) = 4.8, *p* < .001, with larger responses evident in the *Baseline (Guitar)* (*M* = 1.55, *SD* = 0.54) condition than in the *Baseline (Cross)* condition (*M* = 0.9, *SD* = 0.47). To examine whether the pattern of attentional modulation across the two signal types remained consistent when using this more conservative attentional baseline, we calculated new indices of enhancement and suppression using the *attend-Guitars* condition as the baseline (see Figure 5C, right panel) and subjected these indices to the same analyses as reported above. The pattern of enhancement across the two signal types remained intact, with stronger enhancement for the House-selective signal (*M*_chang*e*_ = 1.2, *SD* = 0.87) compared to the bird-selective signal (*M*_chang*e*_ = 0.5, *SD* = 0.51), *F*(1,19) = 15.55, *p* < .001, □^2^ = .201. In both cases enhancement scores departed significantly from zero (one sample *p*_bonf_ values < .01). In contrast, suppression effects differed slightly as a function of which baseline was used: When calculated using *attend-Guitars*, the (paradoxical) slight facilitation effect for the House signal remained numerically positive (*M*_chang*e*_ = 0.14, *SD* = 0.44) but did not differ statistically from zero, *t*(19) = 1.38, p_bonf_ = 0.739), and suppression scores for the bird signal dropped significantly below zero (*M*_chang*e*_ = -0.2, *SD* = 0.19), *t*(19) = -4.77, p_bonf_ < .001 (suppression scores also differed significantly between signal types, *F*(1,19) = 7.59, *p* = .013, □^2^= .203). In other words, attending to houses served to significantly suppress the bird-selective signal when compared to a neutral attentional baseline in which observers had to monitor the images themselves, rather than just a focal part of the display (as in the *attend-Cross* condition).

#### Time Domain

The temporal dynamics of the bird-selective neural response were highly consistent with those observed in Exp 1 (Figure 4D), as was the pattern of attentional enhancement identified by contrasting the Attend Towards and Baseline conditions (where baseline = *attend-Cross*). Just as in Exp 1, explicitly attending to birds facilitated selective neural processing of this category in both an early time window (i.e., between 100-200ms) and, very markedly, during later time window between 320-420ms. Relative to the *attend-Cross* condition, attending to houses attenuated the amplitude of the bird-selective response for a brief period around 100ms post stimulus onset, but resulted in stronger negative amplitudes between 320-420ms. The latter effect differed from what we observed in Exp 1, where attending to the alternate category (Faces) resulted in stronger positive amplitudes for the bird-selective response over a slightly earlier window (∼200-350ms, Figure 4D).

In contrast to the robust and replicable face- and bird-selective responses, the house-selective waveform was less well-formed (see Figure 4C). Under orthogonal task conditions (attend-Cross or attend-Guitars), houses evoked a small initial positivity around 70ms, followed by a prolonged negativity lasting until ∼400ms. A qualitatively similar waveform has been documented previously for houses embedded amid other natural object categories, where the selective response to this category was notably less well-formed than those associated with faces and body parts (Jacques et al. 2016). Attending to houses resulted in a strong positivity during the early stages of the house-selective response, and again during a later time window between 300-400ms. This latter positive component strongly resembled that which was evident in the bird-selective waveform when birds were the attended category. Notably, attending to an alternate category (either birds or guitars) did not serve to dampen house-selective response amplitudes at any timepoint, but rather resulted in stronger long-lasting negativity.

The comparison between the two baseline conditions was also of interest. When the task requires observers to monitor the image stream (but neither critical category is task-relevant, i.e., *attend-Guitars* condition), category-selective signals appear to be facilitated (i.e., larger absolute response amplitudes) relative to when the task requires monitoring of the central fixation cross. For birds, this manifests as a stronger initial positivity and a stronger negativity later on. For Houses, this appears as a stronger initial negativity.

## DISCUSSION

The current study is the first to exploit visual periodicity to dissociate concurrent selective neural signals for different high-level categories contained within the same rapid image stream. We leveraged this powerful ‘interlaced’ periodicities design to capture the spatiotemporal dynamics of attention’s influence on selective visual processing of faces, birds, and houses. Results revealed a qualitatively distinct attentional signature for face-selective visual processing compared to other high-level categories. Where attending to either birds or houses provides a strong boost to the selective neural signals associated with these categories, attentional facilitation of face-selective neural processing appears to be much more modest. Moreover, only the face-selective neural response was actively attenuated by attending to a competing visual category (i.e., minimal or no suppression of the bird- or house-selective response caused by attending to another category). These results underscore the special status that faces hold in the human visual system (Farah 1996; Farah et al. 1998) and highlight the utility of frequency designs as a simple, yet powerful tool for indexing selective neural processing of multiple visual categories contained within the same visual sequence.

### Distinct profiles of attentional enhancement for face and non-face stimuli

The data here establish that monitoring for task-relevant category exemplars amid a large and highly variable set of object images facilitates category-selective neural processing in distinct ways for face vs. non-face categories. When observers in Exp 1 explicitly attended to birds, the selective neural response for this category was 2.63 times as large as the response in the neutral attentional baseline (3.06 times as large in Exp 2). A comparable increase was evident for the house-selective signal in Exp 2, where the selective response in the attend-towards condition was twice as large as the response at baseline. In contrast, attentional facilitation of face-selective visual processing was extremely modest (face-selective response in the attended condition was just 1.17 times as large as the same response at baseline, Exp 1). Inspection of the corresponding category-selective waveforms (i.e., time-domain analysis) revealed distinctions in the temporal unfolding of enhancement effects across the different high-level categories. Although category-based attention facilitated the early stages of selective neural processing for all three categories, the strongest enhancement of the bird- and house-selective waveforms arose much later (i.e., 300-500ms), with no evidence that category-based attention enhanced face-selective visual processing after 200ms.

#### Early facilitation of face-selective neural processing

Early enhancement of selective neural processing for task-relevant faces is consistent with previous reports of a stronger P1-face and a prolonged N1-face when observers actively attend to faces in a rapid image sequence compared to a overlaid fixation point (Quek, Nemrodov*, et al.* 2018). Early attentional enhancement for visual processing of task-relevant (vs. passively viewed) faces has also been found using a standard ERP approach (e.g., increased P1 amplitudes, earlier N1 latencies, Gazzaley et al. 2005; Gazzaley et al. 2008; Zanto et al. 2010). These findings stand apart from studies that have operationalised selective attention as a simple binary (i.e., faces are either the *actively attended* or *actively ignored* category on each trial), where early attentional benefits (i.e., < 200ms post stimulus onset) tend to be evident only under high perceptual load conditions (Lueschow *et al*. 2004; Furey et al. 2006; Sreenivasan et al. 2009). As such, it seems likely that the processing constraints conferred by the fast image presentation rate (6 Hz) and highly variable stimulus set used here improved our chances of observing early enhancement of face-selective processing. Our observation that attending to faces amid a host of other natural categories facilitates the early stages of selective visual processing for this stimulus is also consistent with work linking P1 and N170 amplitudes to categorisation/recognition outcomes for faces (Liu *et al*. 2002). In one MEG study, Liu et al. showed that both M100 and M170 magnitudes are larger in response to degraded face stimuli that were successfully categorised as a faces (albeit only against houses). Notably, only the M170 varied systematically as a function of successful identification (i.e., recognising the individual).

One interesting aspect of our data is that we did not find any evidence for differential attentional enhancement of face-selective signals at left and right occipitotemporal recording sites. Previous work has shown that attending to faces in a rapid visual sequence enhanced face selective processing much more evidently in the non-preferred hemisphere for faces (i.e., the left hemisphere for most participants, +43%) than in the preferred hemisphere (i.e., the right hemisphere for most participants, +3%) (Quek, Nemrodov*, et al.* 2018). In contrast, we found no distinction between the left and right occipitotemporal ROIs in terms of attentional enhancement, with category-based attention boosting the quantified face-selective response to a comparable extent in the left (+22%) and right (+23%) occipitotemporal ROIs. What accounts for this difference is unclear, as the two studies share the same image presentation rate and degree of target/distractor similarity (i.e., both use face targets amid object distractors at a rate of 6 images per second, although the specific stimulus images are different in the two studies). One notable difference between the studies concerns the participants’ task: Where Quek et al. (2018) observers monitored the sequence for instances of *female* faces (just 5 occurrences per sequence), observers in our study had to press the spacebar with their (dominant) right hand in response to all faces. If anything, however, additional motor responses in our *attend-Faces* block would be expected to drive a larger face-selective response in the (contralateral) left hemisphere, which is not the pattern we observe. As such, it seems unlikely that this task difference accounts for the discrepancy in hemispheric attentional differences. Another, more likely, explanation relates to the degree of right-lateralisation in face-selective neural responses in the two participant samples. Where observers in Quek et al. (2018) exhibited predominantly right-lateralised face responses in the neutral attention condition (just 2/15 subjects showed the opposite pattern), our Exp 1 sample showed a much more bilateral distribution of face-selective responses, with 6/20 individuals showing a stronger face-selective response in the left OT ROI in the baseline condition (see Figure S1A in Supplemental material). It is possible that this increased heterogeneity in the spatial distribution of face-selective processing may have masked hemispheric differences in attentional effects. Clearly, more work in this area is needed to determine whether attention truly does exert a differential influence in the left and right face perception networks.

#### Both early and late facilitation for bird- and house-selective neural processing

To the best of our knowledge, the data here are the first attempt to disentangle the timecourse over which category-based attention specifically serves to *i)* enhance and *ii)* suppress selective visual processing for a category other than faces. Over two participant samples, we found that attending to birds reliably enhanced the bird-selective neural signal during an early window between 100-200ms, and again between 300-500ms. A similar pattern of attentional enhancement was evident for house-selective neural processing in Exp 2, with even stronger early enhancement effects. These dynamics are broadly in keeping with the task-relevance effects for object representations that have been documented using neural decoding methods. In one EEG study, object representations elicited by a 2.5 Hz image stream were more dissociable between 200 and 600ms when observers actively monitored the object stream for repeats (Grootswagers et al. 2021), compared to when they attended to a small stream of letters overlaid on the object images. Where these authors speculated that such long-lasting attentional effects might have arisen in part due to the memory-based nature of their 2-back task, the relatively late enhancement of visual processing for birds and houses here (i.e., 300-500ms) cannot be explained a need to hold object identities in mind. Instead, based on the latency and topographical distribution of the positive deflection that arises when these categories are actively attended, we interpret this relatively late enhancement as a P3 component related to attentional selection of target category exemplars (Johnson 1988; Luck and Yard 1995; Hruby and Marsalek 2003). In particular, our data appear consistent with the P3b subcomponent that arises over parietal sites around 300-500ms, which is thought to reflect the cognitive evaluation and categorization of stimuli relevant to the task at hand.

Interestingly, the face-selective waveform also contained a similar positive deflection within the same temporal range, which has in the past been described as a P2-Face (Rossion *et al*. 2015; Jacques *et al*. 2016; Retter and Rossion 2016; Quek, Nemrodov*, et al.* 2018). In our data, this component was evident in both the attend-Faces and baseline conditions, but obliterated when observers actively attended to the alternate category in Exp 1 (birds). This pattern is largely consistent with that observed by Quek et al. (2018), although in their data the late positive component was not entirely suppressed by attending to another category in the sequence. On one hand, this might suggest that the P2-Face reliably observed under orthogonal task conditions (akin to our baseline condition) could comprise a kind of automatic “targetisation” of faces driven by their high salience, which is eliminated when attention is actively directed towards a different target category. On the other hand, however, the spatial topography of the P2-Face is not strictly the same as the more parietal positive scalp distribution typically associated with the P3b, suggesting that their mere coincidence in time should not be reason enough to believe they reflect the same underlying neural generators.

Given that in our paradigm, attending to a category also means responding behaviourally each time that category appears, it is relevant to consider the extent to which our enhancement effects reflect a motoric or premotor contribution. Importantly, our prespecified ROIs do not encompass the central electrodes that typically evidence action preparation signals (e.g., lateralised readiness potentials, or LRPs). Moreover, consequential LRPs should presumably be most evident for stimulus-locked waveforms in the ‘attend-towards’ condition, yet supplemental LRP analyses (Figure S2) show that in 3 out of 4 cases, this prediction was not met. As such, we interpret the enhancement effects here as more likely capturing an increase in selective *visual* processing for the attended category.

### Attentional suppression: Exclusive to face-selective processing?

One contrast to which motoric elements cannot contribute in any way is attentional suppression, as neither the Baseline nor Attend Away condition is associated with a behavioural response at the category-selective frequency. A key finding of the current study is that only face-selective neural processing appeared to be readily suppressed by directing attention towards an alternate high-level category. In Exp 1, monitoring for instances of birds in the image sequence significantly dampened the face-selective signal relative to a neutral attention baseline, yet the reverse was not true. That is, attending to faces had no suppressive effect on the bird-selective signal, and in fact produced a somewhat paradoxical *increase* in the bird-selective response relative to baseline. We suspect this effect is tied to the nature of the participants’ task in this baseline condition: where monitoring the fixation cross demands a narrow, central focus of attention that is much smaller than the images, monitoring for faces necessarily requires attention to the image stream itself. In this way, attending to faces may have in fact created spatial attention based facilitation of bird-selective processing, even though this category was not the focus of task-based attention in that condition. We tested this hypothesis in Exp 2 by including an additional baseline condition where participants monitored the sequence for (non-periodic) instances of a third object category (guitars), which enabled us to quantify both bird- and house-selective processing when the images themselves were attended, but neither category was task-relevant. A direct comparison of the two baseline conditions in Exp 2 confirmed that, as expected, ensuring attention to the image stream itself (by monitoring for guitars) did indeed boost the magnitude of the bird- and house-selective responses, even though participants were not monitoring for instances of those specific categories. As such, we interpret the unexpected increase in category-selective processing observed in the Attend Away condition for birds (Exp 1 & Exp 2) and houses (Exp 2) as a spatial attention facilitation effect.

Faces were the notable exception to this pattern. The face-selective response in the neutral attentional baseline condition in Exp 1 was already very strong, and was clearly suppressed by directing attention towards a non-face category (-8%) (Quek, Nemrodov*, et al.* 2018). Inspection in the time-domain suggested suppression of face-selective processing was largely evidenced after 300ms of visual processing time, enduring until at least 550ms post stimulus onset. Quek, Nemrodov et al. (2018) reported similar effects, however in their case suppression effects were even more pronounced, with strong suppression observed between 200-300ms as well as during the later time window found here. In both studies, monitoring for an alternate high-level category seems to suppress the P2-Face: a face-selective component that is reliably observed using rapid periodic presentation designs (Rossion *et al*. 2015; Quek and Rossion 2017; Quek, Nemrodov*, et al.* 2018). That the P2-Face is suppressed when category-based attention is directed away from faces would support the idea that this component may in part reflect automatic attentional selection of faces as they appear in the image sequence. When visual competition is high (i.e., dynamically changing visual input, fast presentation rate, naturalistic exemplars), perhaps neural architecture that typically prioritises face processing is recruited to serve current task demands (e.g., detect birds).

In contrast to faces, suppression of category-selective processing of non-face stimuli was a more complex picture. As noted above, when quantified against the *attend-Cross* neutral attention baseline, suppression was actually positive for non-face categories, likely reflecting a spatial attention benefit (see above). When we controlled for this spatial attention effect by quantifying suppression against the *attend-Guitars* neutral attention baseline (Exp 2), we found that attending to an alternative category did induce a small but significant suppression of the bird-selective signal (but not the house-selective signal). These subtle suppression effects for non-face stimuli are in line with extant SSVEP studies that have observed that feature based attentional modulation of concurrently presented simple stimuli (e.g., coloured discs rather than natural object categories) is largely explained by enhancement of the attended item’s neural processing, rather than active suppression of known, salient distractors (Forschack et al. 2022). Again, this highlights the unique status of faces as a high-level category that seem to necessitate strong suppression when an alternate category becomes the focus of an observer’s task.

Time-domain analysis provided additional insight into how the temporal unfolding of category-selective processing varied as a function of both the category itself and the focus of category-based attention. Notably, houses were the only category that demonstrated a qualitatively different selective waveform across attention conditions: In effect, where explicitly attending to houses drove both early and late positivity in the house-selective response, attending away from houses (i.e., monitoring for either birds or guitars) seemed to enhance and prolong negativity throughout the house-selective signal. Understanding the combination of factors that accounts for this requires further investigation; clearly, houses are distinct from the other categories of interest here both at a high level (e.g., inanimate, large objects with navigational affordances) but also in terms of low- and mid-level visual features. Building on our results, future studies of attentional enhancement and suppression of high-level visual categories could employ a more comprehensive set of animate and inanimate categories tease out the interplay between category or task-based attention and low-level featural overlap in distractor categories.

### Visual competition in time

Visual periodicity paradigms are ideally suited for revealing the spatiotemporal dynamics of attention’s influence on selective visual processing, since their rapid image presentation rate and highly variable naturalistic image set ensures that objects must compete for visual representation (Desimone and Duncan 1995). Although visual competition is classically associated with spatial clutter (i.e., multiple items present in the same display) (Duncan 1984; Desimone and Duncan 1995; Kaiser et al. 2014), neural competition can also be induced when single items appear in close temporal proximity, as is the case in classic Rapid-Serial Visual Presentation (RSVP) paradigms (Keysers and Perrett 2002; Zivony and Eimer 2020). When images appear in rapid succession, neural representations of individual objects must necessarily overlap. As temporal overlap increases with faster image presentation rates (Grootswagers et al. 2019), both representational strength (i.e., decodability) and category-selective response magnitudes reliably decrease (Retter et al. 2020; Quek et al. 2021). In this way, the current findings suggest that category-based attention ameliorates visual competition induced by temporal clutter, similarly as other higher-order cognitive mechanisms have been shown to do (e.g., multi-object grouping, Quek and Peelen 2020).

Along the same line, our data also suggest that faces are comparatively robust to temporal competition effects, insofar as attentional enhancement for this category was rather modest compared to non-face categories. Moreover, category-selective signals for faces appear to be, at baseline, both larger and more complex than those evoked by non-face categories. In our case, the bird- and house-selective responses reached 33% and 55% of the face-selective response respectively, this is on par with other studies that have observed face-selective signals 2-4 times larger than selective signals associated with houses or body parts (Jacques *et al*. 2016). This would imply that under orthogonal task conditions, faces are less affected by representational competition induced by neighbouring distractors than other high-level categories are. Multiple possibilities exist as to why faces drive such a strong selective neural response amid object distractors, even when they are not explicitly attended to or relevant to current task demands. Face perception is arguably a highly automatized process (for a review, see Palermo and Rhodes 2007) – with some authors arguing that faces can be processed in the absence of attention (Vuilleumier 2000; Reddy *et al*. 2004; Reddy *et al*. 2006; Reddy and Kanwisher 2007; Quek and Finkbeiner 2013). Along this line of thinking, the face-selective signal we measure in the attend-Cross and attend-Bird conditions could reflect a mandatorily-evoked neural response to an ecologically-relevant stimulus arising *outside* the focus of task-based attention (although not necessarily outside the focus of spatial attention, see above). At the same time, it could also be the case that the faces appearing in the sequence are sufficiently salient to briefly *capture* observers’ attention, effectively inducing ‘attentional slippage’ towards these socially relevant stimuli (Lachter et al. 2004; Theeuwes and Van der Stigchel 2006; Crouzet *et al*. 2010; Gluckman and Johnson 2013; Sato and Kawahara 2015). While our study is not designed to distinguish between these and other mechanistic accounts, the data here unequivocally underscore the special status that faces hold in the human visual system as a high-level category that drives a uniquely strong selective neural response even when they are not the stated focus of the observer’s task.

To conclude, the current ‘interlaced’ periodicity-based design provides a simple, yet powerful way to cleanly dissociate selective neural responses associated with different high-level categories contained within the same rapid image stream. The approach offers a significant increase in experimental efficiency by tapping multiple selective neural signals contained within the same sequence, preserving both frequency- and time-domain information. The associated reduction in overall testing duration has a clear advantage in studying visual perception in populations with testing constraints, such as infants, specific patient groups, or individuals in residential care facilities.

## ACKNOWLEDGMENTS

The authors thank Daniela Ferreira Mesquita and Sofia Bonsignore for collecting the data, Fabienne Chetail for experimental programming, and the anonymous reviewers for their helpful comments.

## Footnotes

1 In a sinusoidal contrast manipulation, the first frame in the cycle is displayed at 0% contrast, such that an image will be detectable minimally 16.67ms post stimulus onset (i.e., 20% contrast, see Retter & Rossion, 2016, for comparison between sinewave and squarewave stimulation). This introduces some temporal displacement in the latencies of waveform components shown in the time domain.

## Notes

### Competing Interest Statement

The authors have declared no competing interest.

### Summary of Updates

Minor revisions throughout the text. Additional plot included in the supplemental material.

