## Supplemental Material for "Visual periodicity reveals distinct attentional signatures for face and non-face categories"

SUPPLEMENTAL MATERIALS

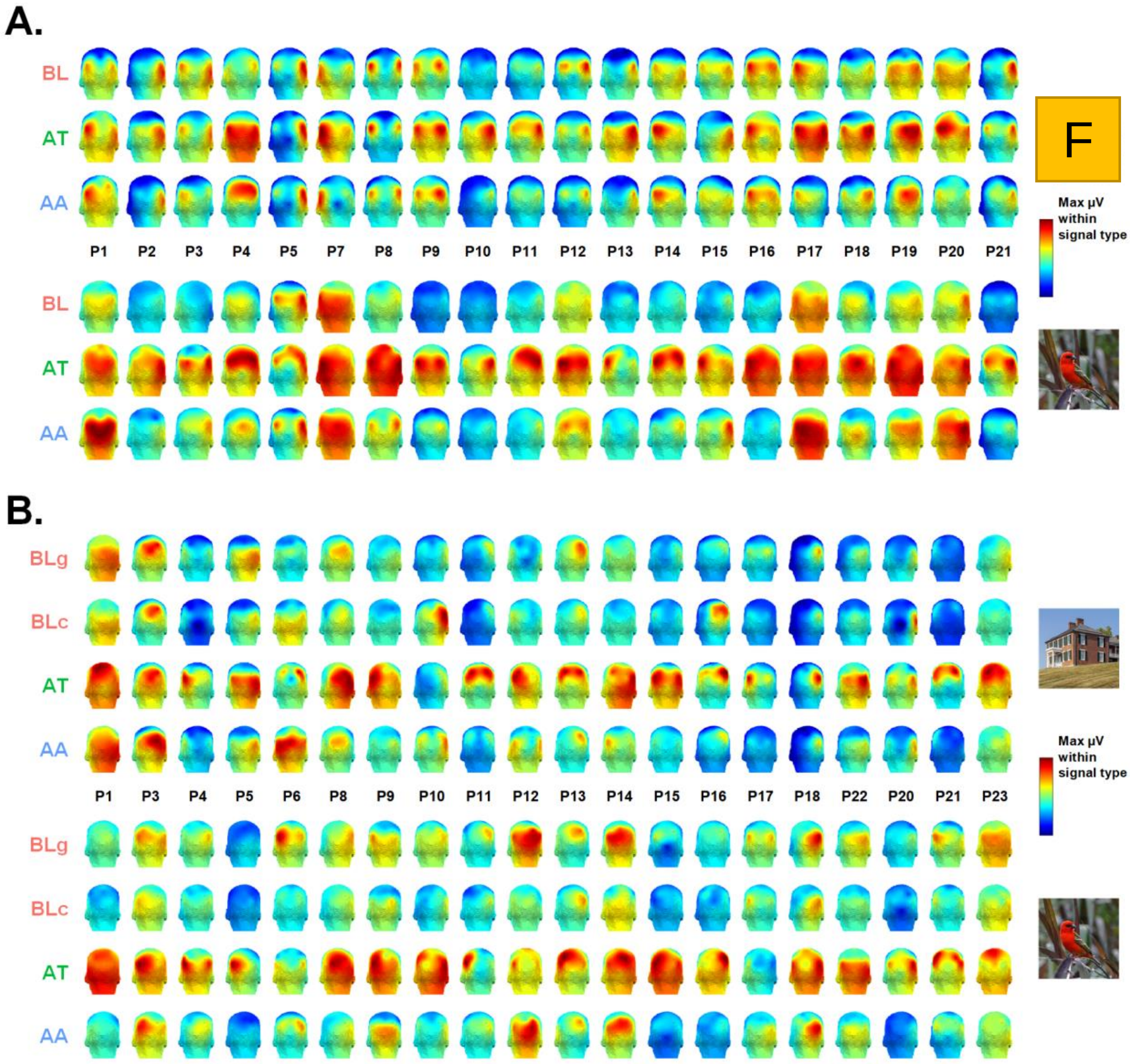


**Figure S1. A)** Individual participant scalp topographies for each category-selective response in each condition of Exp 1, shown as a function of attention condition (BL=baseline; AT = attend towards; AA = attend away). Amplitude ranges are fixed across condition, for each participant individually. **B)** The same for Exp 2 (BLg = baseline (cross); blG = baseline (guitar); AT = attend towards; AA = attend away).

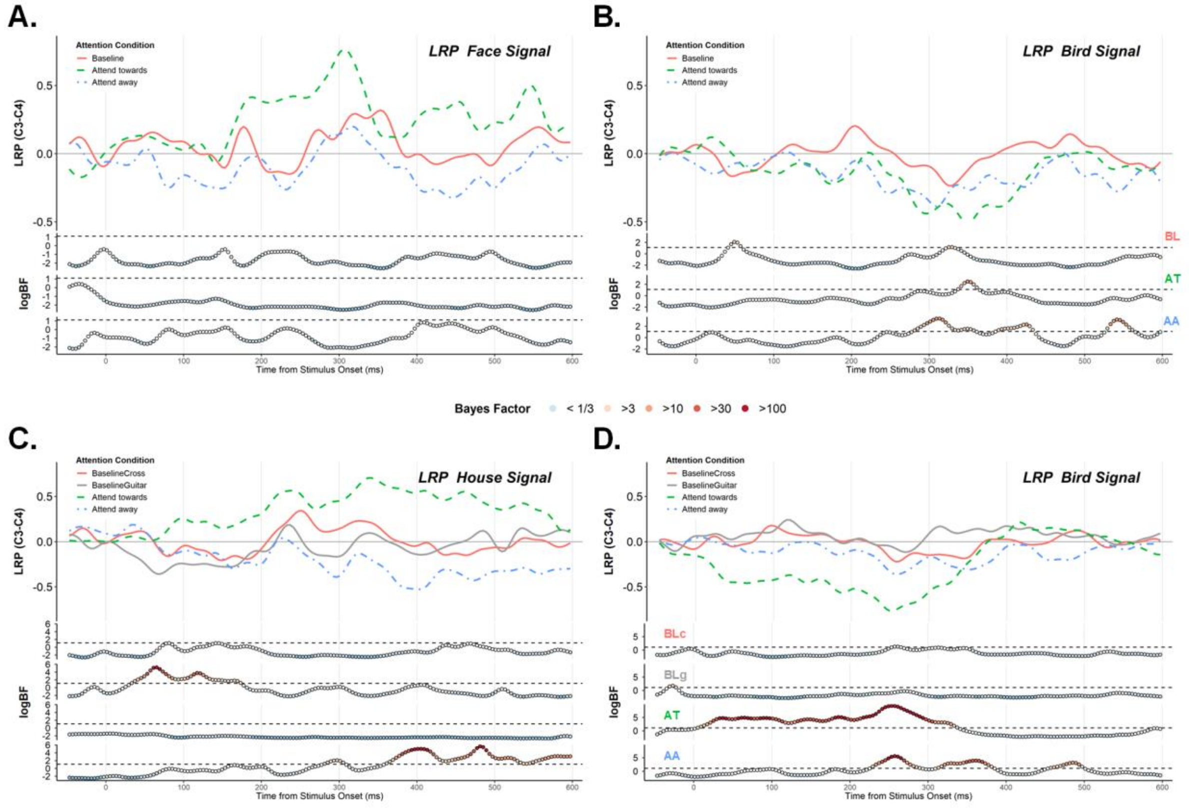


Figure S2. LRPs (electrode C3-C4) calculated using each participant's baseline-corrected conditional mean epoch locked to the onset of A) faces in Exp 1, B) birds in Exp 1, C) houses in Exp 2, and D) birds in Exp 2. Attentional conditions are plotted separately; lower panels show log Bayes Factors testing values < 0 (i.e., a classic negative deflection reflecting motor preparation). We were specifically interested in whether the 'Attend Towards' condition in which participants pressed a button with their right hand each time the critical category appeared on the screen would drive a pronounced LRP. There was no evidence of such a pattern in 3 out of 4 cases.

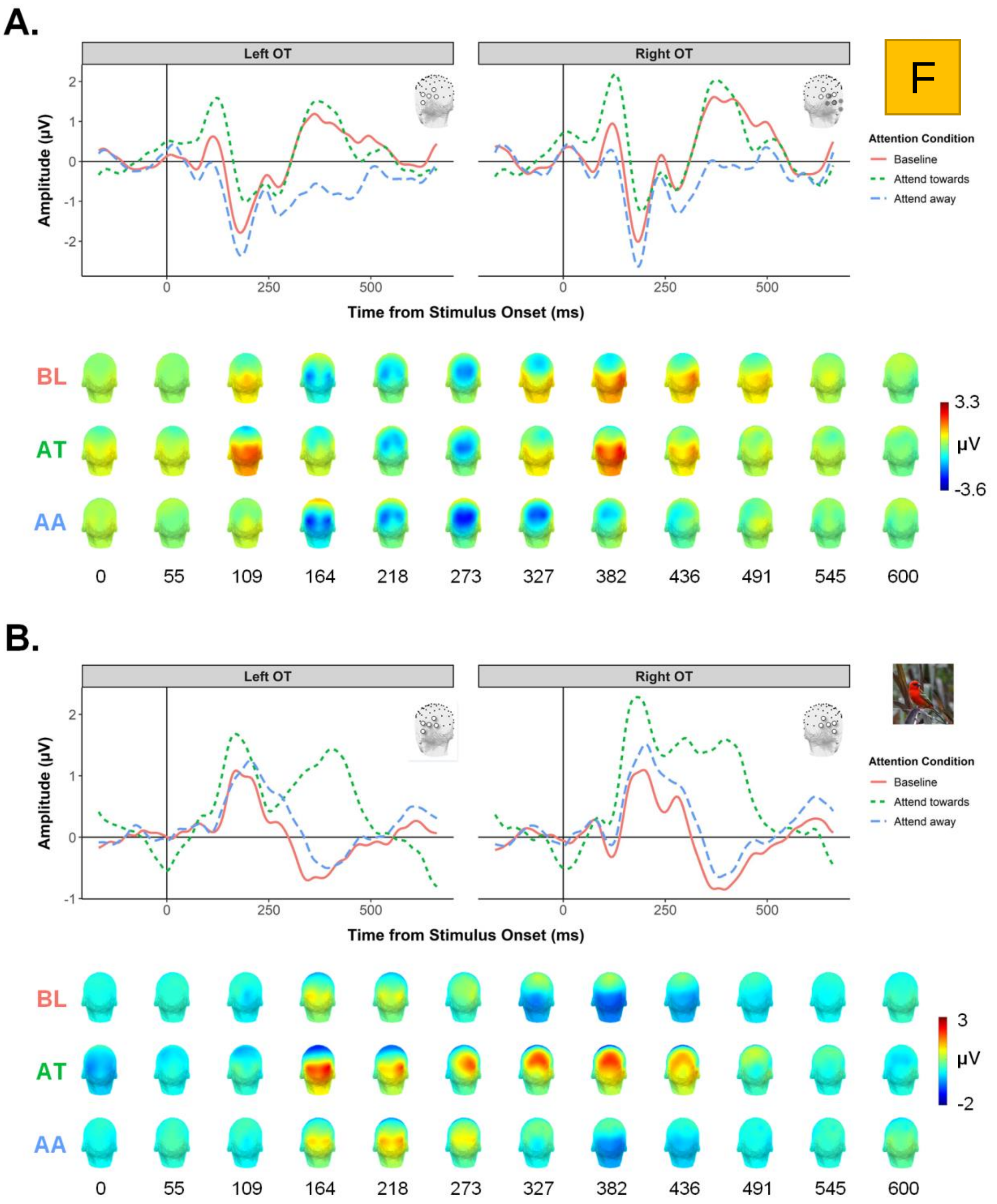


Figure S3. The face-selective (A) and bird-selective responses (B) for Exp 1 shown in the time-domain. Waveforms for each attentional condition are plotted separately for the left and right occipitotemporal ROIs. Group-mean scalp topographies for each condition are shown below each category-selective response for a range of timepoints between 0 and 600ms (BL=baseline; AT = attend towards; AA = attend away).

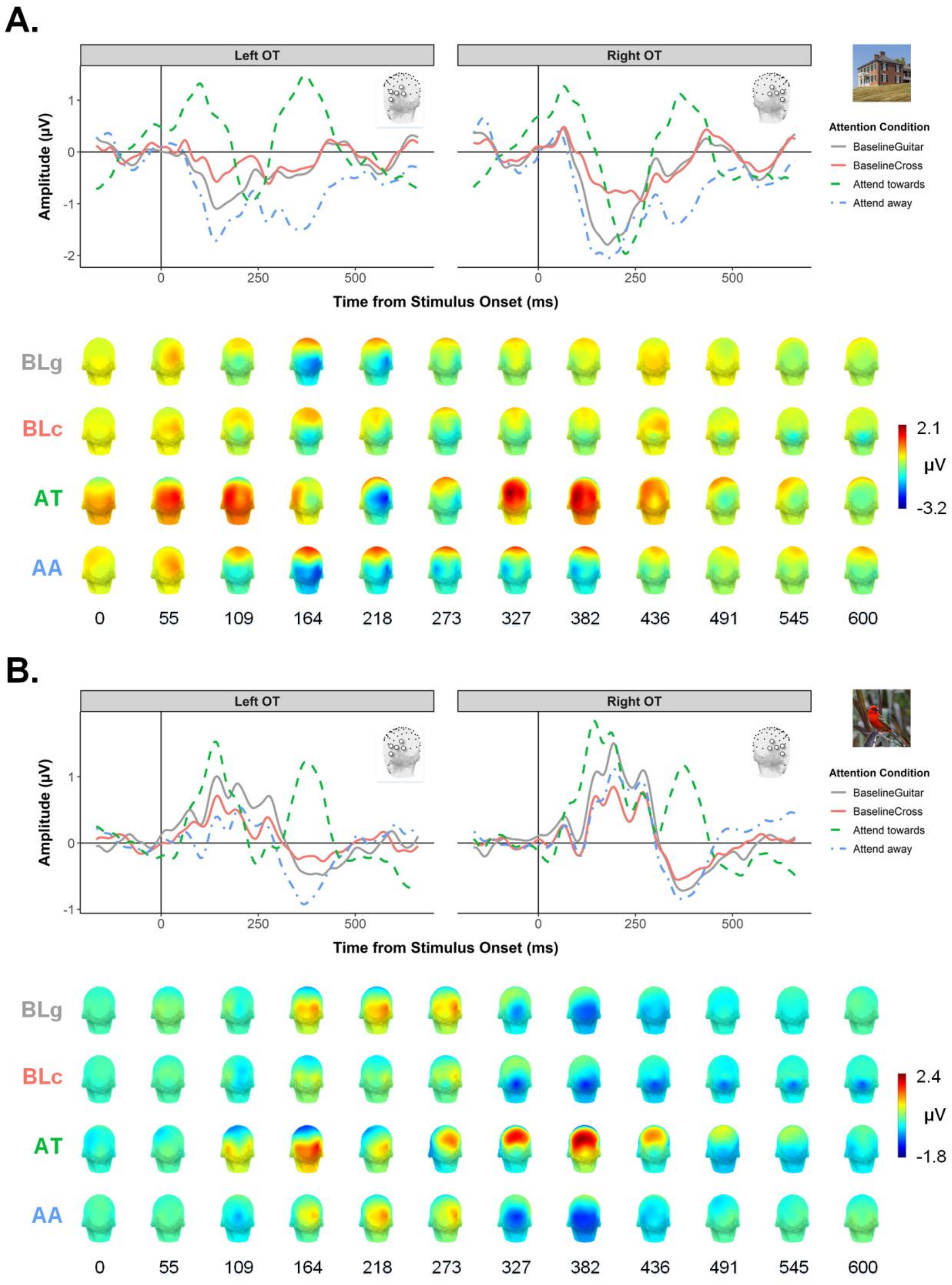


**Figure S4.** The house-selective response (A) and bird-selective response (B) for Exp 2 shown in the time-domain. Waveforms for each attentional condition are plotted separately for the left and right occipitotemporal ROIs. Group-mean scalp topographies for each condition are shown below each category-selective response for a range of timepoints between 0 and 600ms (BLg = baseline guitar; BLc = baseline cross; AT = attend towards; AA = attend away).

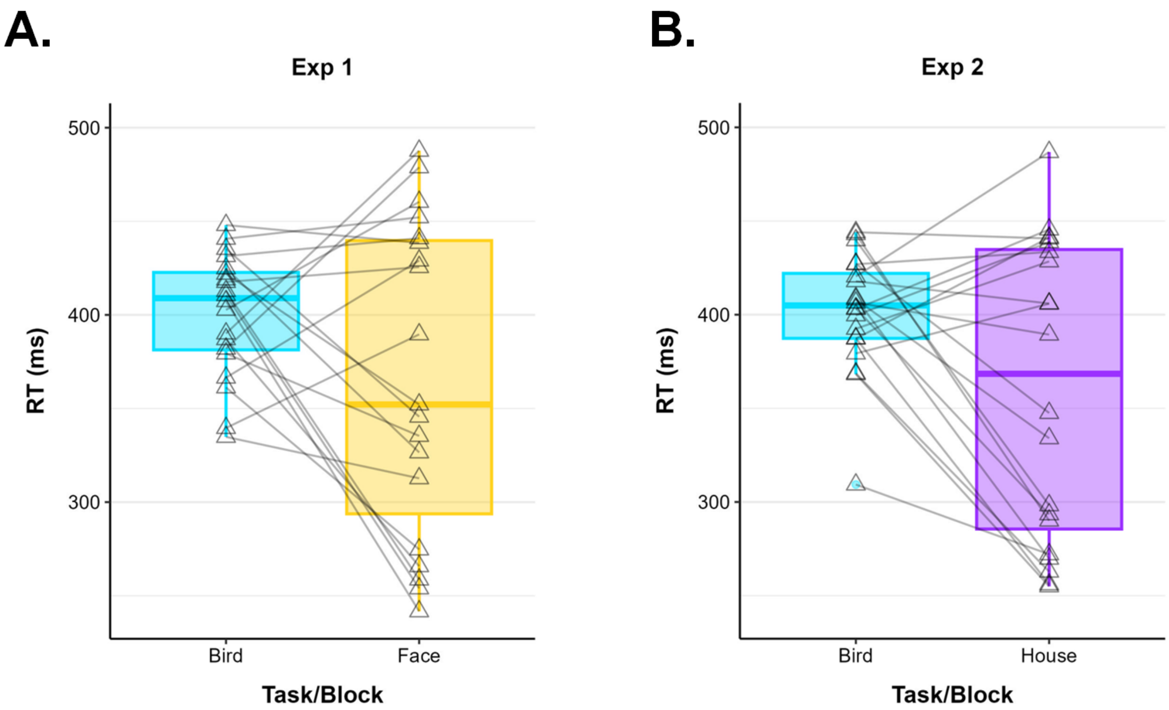


**Figure S5.** Response time (RT) data indexing for participants' detection of the critical categories in (A) Exp 1 (birds/faces) and (B) Exp 2 (birds/houses). Overlaid triangles are individual participants; note that different participant samples were used across the two experiments. In both cases, mean RT was higher for bird-detection responses than for detection responses to the alternate category (Exp 1: $t(19) = 2.01$, $p = 0.059$ ; Exp 2: $t(19) = 2.53$, $p = 0.021$). RTs for both faces and houses also tended to be much more variable than RTs for birds.
